# Mapping the response of primary human chondrocytes to an osteoarthritis-relevant stimulus by single-cell RNA sequencing

**DOI:** 10.64898/2026.09.24.754148

**Authors:** Samantha R Stuppy, Jacqueline Shine, Susan D’Costa, Jorge G Fernandez Davila, Jeremy D Burgess, Adam B Yanke, Richard F Loeser, Douglas H Phanstiel, Brian O Diekman

## Abstract

**Objective:** To determine how fibronectin fragment (Fn-f), a matrikine that induces an osteoarthritis (OA)-like phenotype, reshapes chondrocyte states and transcriptional programs at single-cell resolution.

**Design:** Primary human ankle chondrocytes from two cadaveric donors without OA were treated with PBS or purified recombinant Fn-f for 18 hours and profiled by single-cell RNA sequencing. Transcriptional states were characterized by marker genes and functional enrichment, compared with classifications used for OA chondrocytes, and evaluated for Fn-f-dependent changes in abundance and transcriptional programs.

**Results:** Five transcriptionally distinct populations were identified with features that matched previous classifications of effector, inflammatory, pre-hypertrophic/fibrochondrocyte, reparative, and homeostatic chondrocytes. Fn-f markedly redistributed cells among these states, enriching inflammatory chondrocytes while reducing effector and reparative populations. Within transcriptional states, Fn-f differentially altered matrix/cartilage, inflammatory, and stress/immediate-early programs. These responses did not uniformly correspond to assigned subtype identities, demonstrating that transcriptional programs can be dynamically deployed across chondrocyte states.

**Conclusions:** Controlled perturbations of healthy chondrocytes complement observational OA atlases by resolving stimulus-dependent changes in chondrocyte state abundance and transcriptional programs. Discordance between subtype annotations and dynamic functional programs further highlights the need for harmonized definitions of chondrocytes across phenotypic states.

## Introduction

Osteoarthritis (OA) is the most common form of arthritis and is characterized by progressive cartilage degeneration, extracellular matrix remodeling, and chronic inflammation. Although chondrocytes are the only resident cell type within articular cartilage, single-cell RNA sequencing (scRNA-seq) studies have revealed substantial transcriptional heterogeneity, with chondrocyte populations exhibiting a distinct balance of cellular processes such as matrix synthesis, hypertrophy, and inflammation [1–4]. Across studies, this heterogeneity has been partitioned into an expanding number of proposed chondrocyte states, including effector, pre-hypertrophic, fibrochondrocyte, homeostatic, reparative, and inflammatory populations [5]. Together, these studies have established chondrocyte heterogeneity as an important feature of OA cartilage.

Most OA single-cell studies are observational and focus on a comparison between relatively intact and more damaged regions within OA joints [1–4]. Although essential for defining the landscape of cell states present in diseased tissue, this approach may not be sensitive enough to capture dynamic transcriptional responses that give insight into the drivers of cell state transitions. Mapping single-cell transcriptomics in the context of disease-relevant perturbations provides a complementary strategy by examining chondrocyte responses to defined stimuli under controlled conditions. Gao *et al*. identified divergent transcriptional responses following IL-1β stimulation of human chondrocytes; however, these cells were derived from a single pediatric donor who underwent polydactyl finger excision surgery and were extensively expanded in culture prior to stimulation [6]. Moreover, although the study identified heterogeneous response trajectories, it did not establish how these responses relate to the chondrocyte states that have been observed in human OA tissues.

Our work extends perturbational single-cell approaches to adult primary human articular chondrocytes using fibronectin fragments (Fn-f), which are generated during extracellular matrix degradation in OA joints and initiate signaling pathways through integrin activation [7,8]. Of note, a 42 kDa recombinant form of Fn-f (FN7-10) containing the integrin cell binding domain is a potent matrikine that activates numerous inflammatory and catabolic bulk transcriptomic changes that are characteristic of OA [9,10]. By employing this stimulus in the context of single-cell RNA-seq, we examined both the distribution of transcriptional states and the functional programs operating within those states. Comparison of these experimentally defined populations with chondrocyte subtypes reported across human OA studies was used to evaluate how well current classification frameworks capture dynamic chondrocyte responses.

## Methods

### Human cartilage collection and chondrocyte isolation

In accordance with Institutional Review Board review at Rush University Medical Center and the University of North Carolina at Chapel Hill, human ankle articular cartilage was obtained from two male cadaveric donors aged 56 and 69 years. These donors had no known history of OA or other joint disease. Primary chondrocytes were isolated by sequential pronase and collagenase digestion as previously described [10]. Cells were allowed to recover for two days in high density monolayer culture before 18 hour treatment with serum-free media containing 1 µM purified recombinant fibronectin fragments (Fn-f) or phosphate-buffered saline (PBS) as a vehicle control [10].

### Single-cell RNA sequencing

Single-cell suspensions were prepared according to the Illumina Single Cell 3’ RNA Prep, T10 protocol and sequenced on an Illumina NovaSeq X. Raw sequencing data were processed using Illumina DRAGEN Single Cell RNA software (version 4.4.4) to generate gene-cell count matrices.

### Quality control and clustering

Downstream analyses were performed in Seurat (version 5) using R (version 4.4.0) [11]. Cells were filtered using thresholds of nCount_RNA >2,000 and <25,000, nFeature_RNA >1,000, and mitochondrial transcript content <10%. Gene expression values were log-normalized and the 2,000 most variable genes were identified using FindVariableFeatures(). Linear transformation was performed using ScaleData(). Datasets from both donors and treatment groups were integrated using Harmony (PC = 10) [12]. Cell clusters were identified using the Louvain algorithm at a resolution of 0.2 and visualized using Uniform Manifold Approximation and Projection (UMAP). Cluster marker genes were identified using FindAllMarkers (Wilcoxon rank-sum test) and positive markers were defined as genes significantly enriched within a cluster relative to all remaining cells and were ranked by average log2 fold-change (p < 0.05, logfc.threshold = 0.25, min.pct = 0.1). Clusters were annotated by integrating cluster-specific marker genes with Gene Ontology enrichment analyses and comparison to previously published osteoarthritis single-cell transcriptomic studies.

### Functional enrichment and trajectory analysis

Gene Ontology enrichment was performed using Metascape [13] on cluster marker genes to identify biological processes associated with each transcriptional state. Cell-state trajectories were inferred using Slingshot [14] with cluster 0 (effector chondrocytes) specified as the starting population. Pseudotime trajectories were visualized on the integrated UMAP.

### Transcriptional program analysis

Gene sets representing three transcriptional programs were evaluated for changes in response to treatment: matrix/cartilage (COL2A1, ACAN, COMP, FRZB, CHAD, HAPLN1), inflammatory (CXCL8, IL1B, IL6, PTGS2, CXCL1, CXCL2, CXCL3), and stress/immediate-early (JUN, FOSB, ATF3, DDIT3, MYC, HSPA1A). Module scores were calculated at the individual-cell level using Seurat AddModuleScore() on log-normalized RNA expression with default control-gene parameters and a fixed random seed (123). Scores were summarized by donor, transcriptional state, and treatment condition, and donor-level means were used to compare program activity between PBS- and Fn-f-treated cells. To visualize the genes contributing to each program, normalized expression was averaged for each gene within each transcriptional state and treatment condition and subsequently scaled across state-treatment groups on a gene-wise basis.

## Results

Primary human ankle chondrocytes from two donors were treated with PBS or Fn-f for 18 hours prior to scRNA-seq (Figure 1A). After filtering, 11,297 PBS-treated and 6,171 Fn-f-treated cells from Donor 1 and 10,548 PBS-treated and 4,729 Fn-f-treated cells from Donor 2 were retained for downstream analysis (Supplemental Figure 1). Five transcriptionally distinct chondrocyte populations were identified. Trajectory inference further suggested relationships among these states, with transcriptional paths extending from effector chondrocytes (EC) toward reparative (RepC), pre-hypertrophic/fibrochondrocyte-like (preHTC/FC), and then diverging towards the homeostatic (HomC) or inflammatory (InfC) states (Figure 1B).

**Figure 1.**
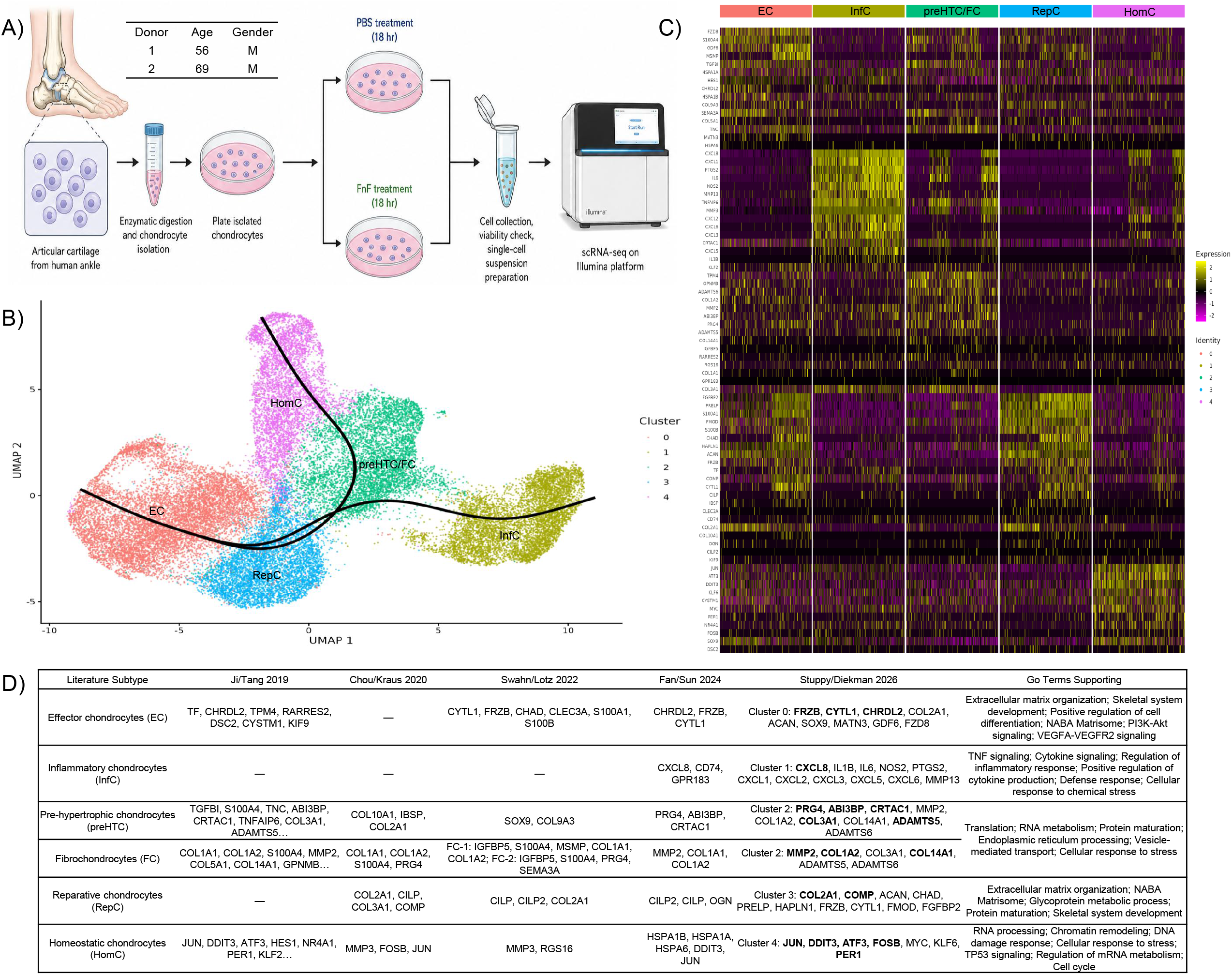
Identification and characterization of transcriptional chondrocyte states in a controlled Fn-f perturbation model. (A) Experimental design. Primary human ankle articular chondrocytes were isolated from two male cadaveric donors, cultured for two days, serum-starved, and treated with phosphate-buffered saline (PBS) or fibronectin fragments (Fn-f) for 18 hours prior to single-cell RNA sequencing. Donor age and Collins cartilage grades are shown. (B) UMAP of integrated chondrocytes showing five transcriptionally defined populations annotated as effector chondrocytes (EC), inflammatory chondrocytes (InfC), pre-hypertrophic/fibrochondrocyte-like chondrocytes (preHTC/FC), reparative chondrocytes (RepC), and homeostatic chondrocytes (HomC). Black curves indicate transcriptional trajectories inferred using Slingshot with EC specified as the starting population. (C) Heatmap showing expression of chondrocyte subtype-associated markers compiled from the published human OA single-cell studies summarized in panel D. Genes are grouped according to their literature-associated subtype. (D) Cross-study comparison of chondrocyte subtype definitions. Representative markers reported for effector, pre-hypertrophic, fibrochondrocyte, homeostatic, reparative, and inflammatory chondrocytes are compared with markers identified in the corresponding populations in the present study. Bold genes indicate overlap with prior studies. Gene Ontology terms enriched among cluster markers provide additional functional support for state annotation. Dashes indicate subtypes not described in the corresponding study.

To relate these populations to chondrocyte states described in human OA, we examined expression of subtype-associated markers reported across published single-cell studies (Figure 1C). Literature-derived markers were sufficient to match the cell states observed in this study, despite some genes showing high expression in several of our defined clusters. Integration of these expression patterns with Gene Ontology enrichment and cross-study comparison supported annotation of the five observed clusters as EC, InfC, preHTC/FC, RepC, and HomC (Figure 1D). Correspondence was relatively clear for InfC, whereas other populations shared features across published subtype definitions. Most notably, cluster 2 exhibited markers attributed to both pre-hypertrophic and fibrochondrocyte populations. Together, these analyses demonstrate broad correspondence with chondrocyte states identified in human OA while highlighting overlap in the markers and terminology used to define them across studies.

Stratification by treatment revealed marked differences in the distribution of PBS- and Fn-f-treated cells across the transcriptional landscape (Figure 2A). EC and RepC were predominantly represented under PBS conditions, whereas Fn-f-treated cells overwhelmingly populated InfC and were also prominent within preHTC/FC. Quantification of state composition confirmed these differences, with InfC comprising the largest proportion of Fn-f-treated cells while being nearly absent under PBS conditions (Figure 2B).

**Figure 2.**
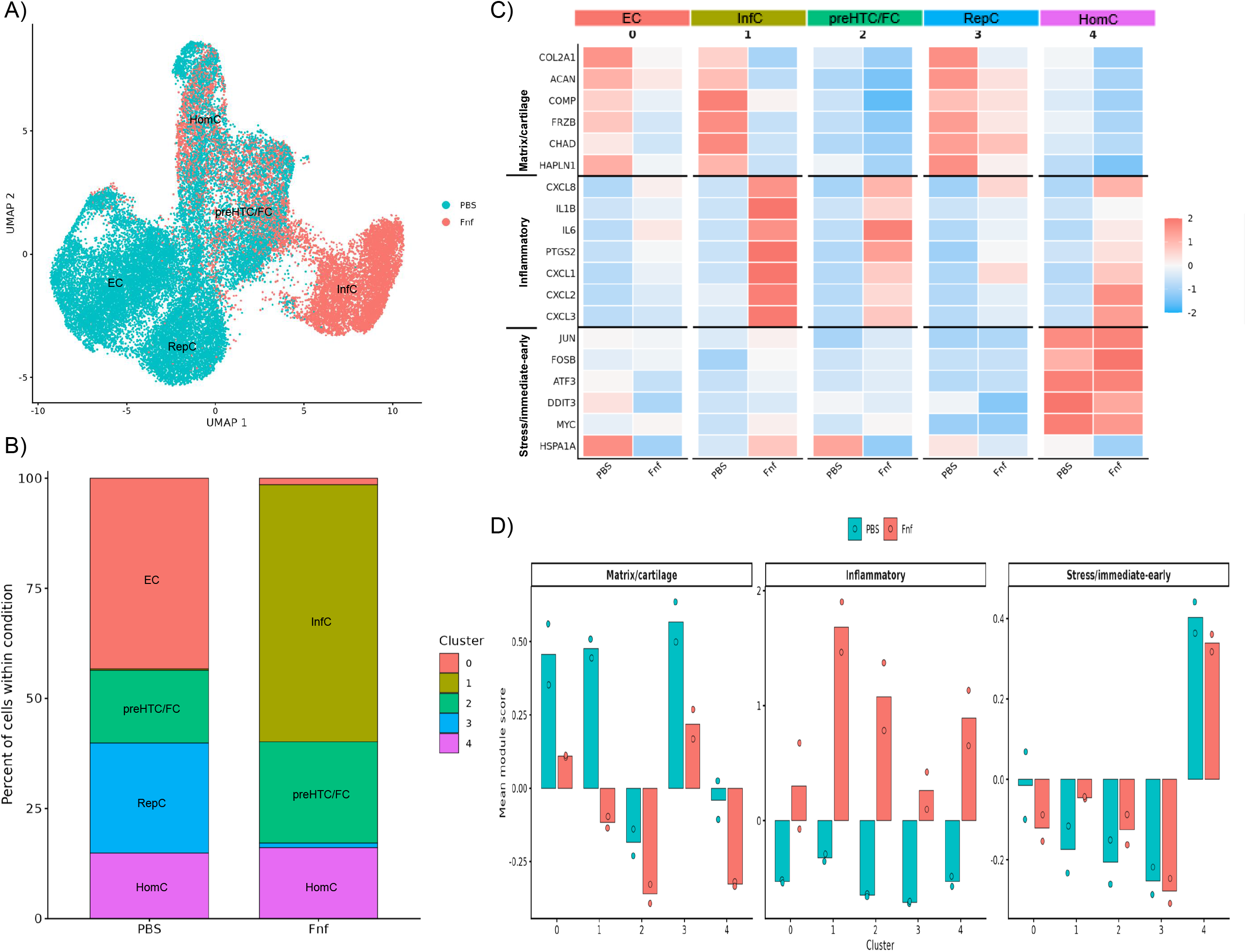
Fn-f induces state-dependent remodeling of chondrocyte composition and transcriptional programs. (A) Integrated UMAP colored by treatment condition (PBS or Fn-f), with transcriptional state annotations shown for reference. (B) Relative abundance of EC, InfC, preHTC/FC, RepC, and HomC within PBS- and Fn-f-treated cells. Bars represent the percentage of cells assigned to each state within treatment condition. (C) Heatmap showing expression of genes representing matrix/cartilage, inflammatory, and stress/immediate-early transcriptional programs across chondrocyte states and treatment conditions. Salmon and blue indicate relatively higher and lower expression, respectively. (D) Matrix/cartilage, inflammatory, and stress/immediate-early module scores across chondrocyte states and treatment conditions. Points represent donor-level mean scores and bars represent the mean across donors.

We next examined whether Fn-f altered functional transcriptional programs within each state (Figure 2C). Matrix/cartilage genes generally showed lower expression following Fn-f treatment, although the magnitude of this response varied among states. Inflammatory genes showed the strongest coordinated induction in InfC but were also induced across other populations, particularly preHTC/FC. HomC exhibited a distinct immediate-early/stress signature, with prominent expression of JUN, FOSB, ATF3, DDIT3, and MYC that was evident even under PBS conditions.

Module scoring provided a program-level summary of these responses across donors (Figure 2D). Fn-f was associated with lower matrix/cartilage scores and higher inflammatory scores across all five states, with the largest inflammatory response in InfC. In contrast, stress/immediate-early scores were highest in HomC and generally lower following Fn-f treatment. Thus, Fn-f altered both the relative abundance of chondrocyte states and the transcriptional programs expressed within them. Notably, these responses were not restricted to states with corresponding functional labels; for example, inflammatory program scores increased across all five states rather than exclusively in InfC.

## Discussion

Fn-f, a matrix degradation product that acts as a “matrikine”, altered both the distribution of transcriptionally defined chondrocyte states and the functional programs operating within those states. Fn-f produced a particularly striking shift toward the InfC state, consistent with the established inflammatory and catabolic effects of this OA model. Fn-f increased the inflammatory module score across all five populations, while matrix/cartilage programs were generally reduced. Thus, expression of an inflammatory program was not synonymous with inflammatory cell-state identity. A cell could retain the broader transcriptional features that placed it within EC, preHTC/FC, RepC, or HomC while simultaneously exhibiting an increased inflammatory program following Fn-f stimulation. PreHTC/FC exhibited substantial inflammatory activation despite retaining a transcriptional profile most consistent with matrix-remodeling, pre-hypertrophic, and fibrocartilage-associated populations. The HomC population exhibits a strong immediate-early/stress program that is largely independent of Fn-f exposure, aligning more closely with the transcriptional profile. This distinction between transcriptional state and functional program is difficult to resolve from static comparisons of human tissue and illustrates a particular advantage of controlled perturbational approaches. These observations caution against interpreting subtype names as fixed descriptions of cellular function. Rather, transcriptionally defined chondrocyte states appear capable of engaging context-dependent programs that cross conventional subtype boundaries.

Our study also highlights challenges in current OA chondrocyte classification. Although the five identified populations broadly resembled chondrocyte states reported in human OA cartilage, cross-study comparison revealed that these states are often defined by different sets of marker genes and our clusters had some features that aligned with multiple states. The strongest overlap was the cluster that showed a strong identity of pre-hypertrophic and fibrochondrocyte-associated markers. Such discrepancies do not necessarily indicate that individual classification systems are incorrect; rather, they may reflect biological continuum among chondrocyte states, differences in tissue source and disease context, analytical approaches, and classification based partly on context-dependent functional programs. Greater harmonization of chondrocyte nomenclature and molecular definitions would facilitate comparisons across OA single-cell studies. Future classification frameworks may particularly benefit from distinguishing relatively stable aspects of cell-state identity from dynamic programs such as inflammation, matrix synthesis and remodeling, and cellular stress.

Several limitations should be considered. Chondrocytes were examined following isolation and short-term monolayer culture, which does not reproduce the native cartilage environment, including extracellular matrix architecture, mechanical loading, or interactions with other joint-resident cell types. The study included only two donors, and while this limits our ability to characterize inter-individual variability in Fn-f responses, a previous study with 101 donors demonstrated that consistency across donors is a strength of the model system [10].

Ankle cartilage may also differ from the knee cartilage which has been examined in many published OA single-cell studies. Prior studies found that ankle chondrocytes isolated from the tali respond similarly to Fn-f as knee chondrocytes isolated from the femur [9,15]. Finally, the focused transcriptional programs evaluated here were selected to interrogate prominent biological features of the identified states and do not represent a comprehensive framework for chondrocyte function.

Together, these findings demonstrate the value of incorporating disease-relevant perturbations alongside observational human tissue atlases to interrogate chondrocyte biology. Perturbational scRNA-seq can reveal both changes in the representation of transcriptionally defined states and changes in the functional programs operating within those states. Applying such approaches across donors, stimuli, and experimental systems, together with efforts to harmonize chondrocyte nomenclature, may help distinguish reproducible cell states from context-dependent transcriptional programs and provide a more consistent and functionally informative framework for understanding chondrocyte heterogeneity in OA.

## Acknowledgments

The authors gratefully thank the families of tissue donors affiliated with Gift of Hope Organ and Tissue Donor Network, Rush University, and Mrs. Arnavaz Hakimiyan for their support in tissue procurement. The authors thank Illumina for providing materials through a single-cell RNA sequencing workshop.

## Author Contributions

Conception and design: SRS, BOD

Analysis and interpretation of the data: SRS, RFL, DHP, BOD

Drafting of the article: SRS

Critical revision of the article for important intellectual content: JS, SDC, JGFD, JDB, ABY, RFL, DHP, BOD

Provision of study materials or patients: ABY

Statistical expertise: SRS, DHP

Obtaining of funding: BOD

Administrative, technical, or logistic support: JS, SDC, JGFD, JDB

Collection and assembly of data: SRS, JS, SDC, JGFD, JDB

Final approval of the article: All authors

## Declaration of Generative AI and AI-assisted Technologies in the Writing Process

During the preparation of this work, the authors used OpenAI ChatGPT to assist with condensing the manuscript to meet the journal word limit. After using this tool, the authors reviewed and edited the content as needed and take full responsibility for the content of the publication.

---

**Supplementary Figure S1.**
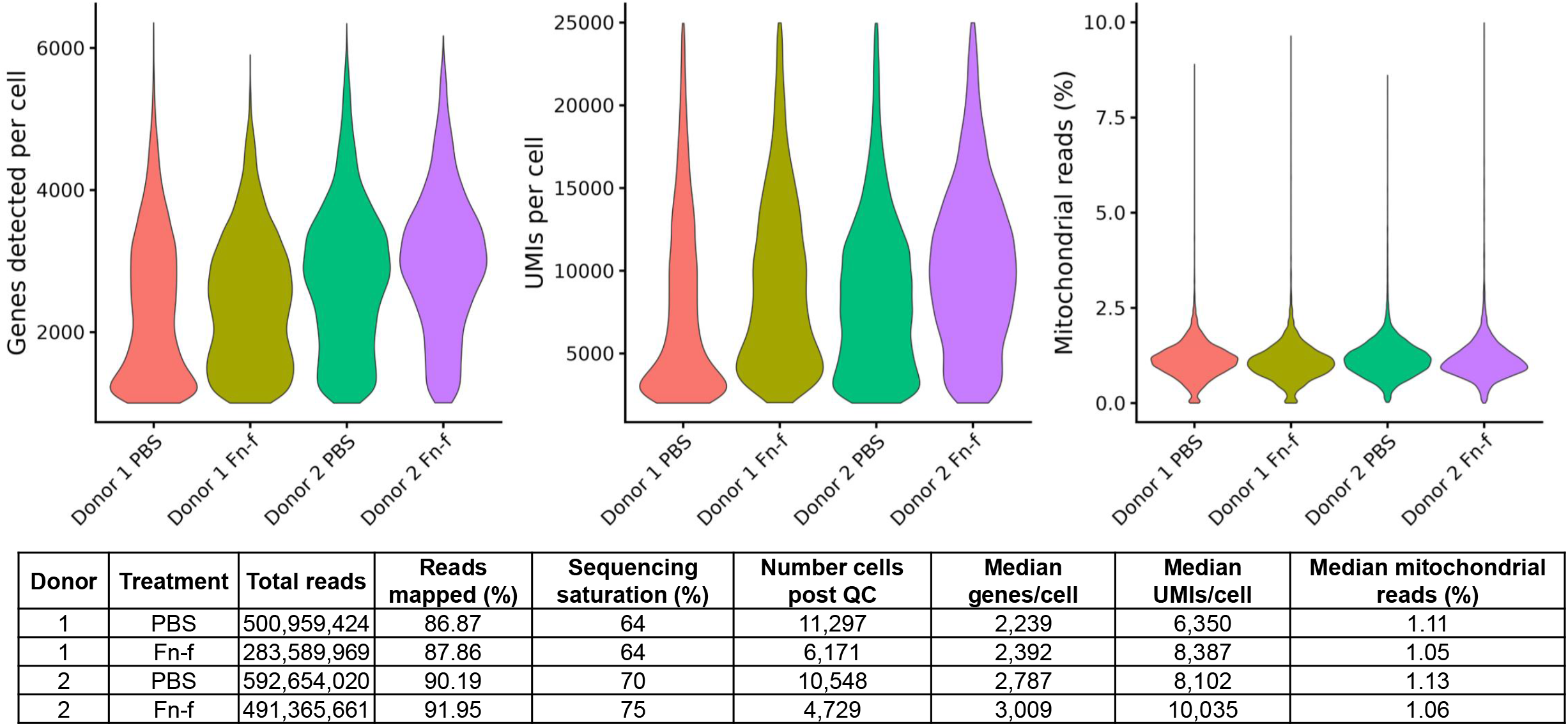
Sequencing and quality-control metrics for single-cell RNA-seq samples. Violin plots show the distributions of genes detected per cell, UMI counts per cell, and mitochondrial RNA percentage among cells retained following quality-control filtering. The table summarizes sequencing and cell-level quality-control metrics for each donor and treatment condition. Total reads, mapped reads, and sequencing saturation were obtained from DRAGEN Single Cell RNA processing metrics; cell counts and cell-level metrics reflect cells retained following Seurat filtering.

